# Sensitization of knee-innervating sensory neurons by tumor necrosis factor-α activated fibroblast-like synoviocytes: an in vitro, co-culture model of inflammatory pain

**DOI:** 10.1101/791251

**Authors:** Sampurna Chakrabarti, Zoe Hore, Luke A. Pattison, Sylvine Lalnunhlimi, Charity N. Bhebhe, Gerard Callejo, David C. Bulmer, Leonie S. Taams, Franziska Denk, Ewan St. John Smith

**Affiliations:** Department of Pharmacology, University of Cambridge; Wolfson Centre for Age-Related Diseases, Institute of Psychiatry, Psychology & Neuroscience; Centre for Inflammation Biology and Cancer Immunology, Department of Inflammation Biology, School of Immunology & Microbial Sciences, King’s College London

**Keywords:** synoviocytes, sensory neurons, pain, inflammation, tumor necrosis factor, complete Freund’s adjuvant, knee

## Abstract

Pain is a principal contributor to the global burden of arthritis with peripheral sensitization being a major cause of arthritis-related pain. Within the knee joint, distal endings of dorsal root ganglion neurons (knee neurons) interact with fibroblast-like synoviocytes (FLS) and the inflammatory mediators they secrete, which are thought to promote peripheral sensitization. Correspondingly, RNA-sequencing has demonstrated detectable levels of pro-inflammatory genes in FLS derived from arthritic patients. This study confirms that stimulation with tumor necrosis factor (TNF-α), results in expression of pro-inflammatory genes in mouse and human FLS (derived from OA and RA patients), as well as increased secretion of cytokines from mouse TNF-α stimulated FLS (TNF-FLS). Electrophysiological recordings from retrograde labelled knee neurons co-cultured with TNF-FLS, or supernatant derived from TNF-FLS, revealed a depolarized resting membrane potential, increased spontaneous action potential firing and enhanced TRPV1 function, all consistent with a role for FLS in mediating the sensitization of pain-sensing nerves in arthritis. Therefore, data from this study demonstrate the ability of FLS activated by TNF-α to promote neuronal sensitization, results that highlight the importance of both non-neuronal and neuronal cells to the development of pain in arthritis.

## Introduction

Joint inflammation and pain are the major clinical symptoms of arthritis. Inflammation is part of the body’s immune response to tissue damage and involves multiple cell types, including leukocytes and synoviocytes in synovial joints. These non-neuronal cells are either in direct contact with, or in close proximity to, the distal endings of joint-innervating dorsal root ganglion (DRG) sensory neurons, and this interaction is thought to cause peripheral sensitization and development of inflammatory pain.

During inflammatory diseases such as rheumatoid arthritis (RA), the affected joints undergo hyperplasia due to leukocyte infiltration and fibroblast-like synoviocyte (FLS) proliferation. FLS are key effectors in RA as they become active upon stimulation by inflammatory cytokines (released by macrophage-like synoviocytes, and T-lymphocytes) and secrete matrix metalloproteases (MMP) that cause joint destruction [9]. Correspondingly, single-cell transcriptional analysis of human RA-FLS, has identified distinct “destructive” and “inflammatory” FLS subgroups [17]. In addition, FLS can also support and maintain the ongoing inflammation in arthritic joints by secreting pro-inflammatory mediators themselves [9]. To better understand the inflammatory phenotype of FLS, several studies have used cytokine stimulation (e.g. interleukin 1β, IL-1β, or tumor necrosis factor-α, TNF-α) to robustly induce inflammation [14,27,30,40]. “Inflamed” FLS have been utilized in co-culture systems to dissect the complex interaction between FLS and other cell types in the joint environment. Co-culture of human RA-FLS with T-lymphocytes [7,11,37,67] or macrophages/monocytes [6,8,15] has been found to increase the concentration of prostaglandin (PG) E_2_, IL-6, IL-8, MMP-1 and MMP-3 in the culture medium. Furthermore, upregulation of these pro-inflammatory mediators in culture supernatants was inhibited by anti-TNF-α [11,55] and anti-IL-6 antibodies [55].

However, most co-culture studies have focused on immune interactions during joint inflammation and therefore, the communication between neurons and synoviocytes is much less understood. FLS have been hypothesized to enhance nociceptive responses [64] and alter the bio-mechanical microenvironment of DRG neurons [31]. FLS derived from chronic antigen-induced arthritic (AIA) rats were observed to increase the expression of receptors associated with nociception, namely neurokinin 1, bradykinin 2 and transient receptor potential vanilloid 1 (TRPV1) in DRG neurons when co-cultured [4]. However, this study did not functionally assess modulation of DRG neuron excitability, which is a key mechanism of peripheral sensitization and hence pain. We recently showed that human osteoarthritic (OA) synovial fluid (a lubricating fluid largely secreted by FLS [5]) can cause hyperexcitability of murine sensory neurons and increase TRPV1 function [12].

To establish a suitable pro-inflammatory phenotype for our cultures, we utilized primary FLS derived from mice undergoing acute, unilateral complete Freund’s adjuvant (CFA)-induced knee inflammation, which can produce neuronal hyperexcitability [13]. We also tested primary mouse knee-derived FLS treated with TNF-α, one of the main cytokines upregulated in CFA-injected mouse tissues [65] and in inflammatory arthritis [53], and validated our results in FLS derived from human OA and RA patients. The pro-inflammatory phenotype of FLS was established to test the hypothesis that after induction of inflammation, mouse knee-derived FLS will increase excitability and TRP function of knee-innervating DRG neurons (knee neurons) in an FLS-DRG neuron co-culture system.

## Methods

### Animals

All mice used in this study were 6-12 week old C57BL/6J mice (Envigo) randomly allocated to each experiment. Unless otherwise stated, female mice were used since in humans, females are at a higher risk for arthritic pain [28,51,62]. Mice were housed in groups of up to 5 in a temperature controlled (21 °C) room with appropriate bedding materials, a red shelter and enrichment. They were on a 12-hour/light dark cycle with food and water available *ad libitum*. Experiments in this study were regulated under the Animals (Scientific Procedure) Act 1986, Amendment Regulations 2012. All protocols were approved by a UK Home Office project license granted to Dr Ewan St. John Smith (P7EBFC1B1) and reviewed by the University of Cambridge Animal Welfare and Ethical Review Body.

### Knee injections

Under anesthesia (100 mg/kg ketamine and 10 mg/kg xylazine, intra-peritoneally) mice were injected intra-articularly through the patellar tendon into each knee with the retrograde tracer, fast blue (FB, 1.5 μl 2% in 0.9% saline, Polysciences) or into the left knee with 7.5 μl CFA (10 mg/ml, Chondrex). Vernier’s calipers were used to measure knee width (as before [13]) pre- and 24-hour post-CFA injection.

### Isolation and culture of FLS

24-hours after CFA injection into the knee, mice were killed by cervical dislocation and decapitation. Knee joints were exposed by removing the skin, then the quadriceps muscles were resected in the middle and pulled distally to expose the patellae. Patellae were then collected by cutting through the surrounding ligaments, as described before [25], in phosphate-buffered saline (PBS) and then transferred into one well of a 24-well plate with FLS culture media containing: Dulbecco’s Modified Eagle Medium F-12 Nutrient Mixture (Ham) (Life Technologies), 25% fetal bovine serum (Sigma), 2 mM glutamine (Sigma) and 100 mg/ml penicillin/streptomycin (Life Technologies). Cells took approximately 10 days to grow to 70% confluency; medium was changed every other day. For P1, FLS were trypsinized with 1% trypsin (Sigma), re-suspended in FLS culture media and transferred into two wells of a 6-well plate. FLS from two animals were combined at P2. For subsequent passages, FLS were transferred into 60 mm dishes. Contralateral (Contra) and CFA-injected knees/cells (Ipsi) were kept separate at all stages. The cells were maintained in a humidified, 37 °C, 5% CO_2_ incubator. FLS were also cultured until P5 from mice that had not undergone any knee CFA injection (control). A random selection of these dishes, from three separate biological replicates, was incubated for 24 to 48-hours (as per experimental design) in culture medium with recombinant mouse TNF-α (10 ng/ml from a stock solution of 100 μg/ml made up in 0.2 % bovine serum albumin and sterile PBS, R&D systems, aa-80325) to stimulate release of inflammatory mediators (TNF-FLS) [29].

### Culture of Raw 264.7 cells

Raw 264.7 cells (EACC) were cultured in medium containing Dulbecco’s Modified Eagle Medium F-12 Nutrient Mixture (Ham) (Life Technologies), 10% fetal bovine serum (Sigma), 2 mM glutamine (Sigma) and 100 mg/ml penicillin/streptomycin (Life Technologies). Cells were maintained in a humidified 37 °C, 5% CO_2_ incubator. This macrophage-like cell line [59] was used to assess the level of macrophage contamination in FLS cultures.

### RNA extraction and reverse transcriptase – quantitative/ polymerase chain reaction (RT-q/PCR) of mouse FLS

For all conditions, RNA was extracted from two 60 mm dishes (Thermo Fisher) of FLS (various passages) and from one T-25 flask (Greiner Bio-one) of Raw 264.7 cells at P3 using the RNeasy Mini Kit (Qiagen). 500 ng of the extracted RNA was used to synthesize cDNA using a High Capacity cDNA RT kit (Applied Biosystems) following the manufacturer’s guidelines, using a T100 Thermal Cycler (Bio-Rad). The resultant cDNA was diluted to a 1:5 ratio with nuclease free water and quantitative PCR (qPCR) was performed using a StepOnePlus Real Time PCR system, following the manufacturer’s guidelines on settings (Applied Biosystems) using TaqMan probes (Thermo Fisher) (Supplementary Table 1 listing genes of interest analyzed in this study). The fluorescence intensity of samples was captured during the last minute of each cycle. All reactions were run in triplicate with appropriate negative controls with water containing no cDNA.

For RT-qPCR reactions, data were obtained as Ct values (the cycle number at which fluorescent signals emitted by the TaqMan probe crossed a threshold value). Only Ct values below 35 were analyzed to determine ΔCt values by subtracting the Ct of *18S* ribosomal RNA from the Ct of target gene [41]. ΔCt values of target genes were subtracted from average ΔCt values of their controls to calculate ΔΔCt, followed by 2^(-ΔΔCt) to calculate fold change [39].

FLS gene expression was also assessed by RT-PCR. DreamTaq Polymerase (Thermo Fisher) was used to amplify a section within the open reading frames of various genes from 5 ng template cDNA. The sequences of designed oligonucleotides (Sigma) are listed in Supplementary Table 1. Negative controls (using water/RNA as the template) were performed for each biological sample with a randomly selected primer pairing. PCR products were resolved on 2% agarose containing 1X GelRed Nucleic Acid Stain (Biotum) and imaged with a GeneFlash Gel Documentation System (Syngene). A randomly selected subset of reactions was repeated with 2.5 ng template cDNA to ensure reproducibility. Densitometry analyses were performed using ImageJ Software (NIH) where relative expression was determined by dividing the band intensity of each gene by that of the housekeeping gene, *18S* ribosomal RNA, for each biological replicate.

### Cytokine antibody array

Before RNA extraction, 2 ml of culture media from P5 Contra, Ipsi, control and TNF-FLS (48-hour) were collected and stored at −80 °C until use. Culture medium was pooled from three cultures for each of the four conditions and assayed (undiluted) for the presence of 40 inflammatory mediators using Mouse Inflammatory Antibody Array Membranes (ab133999, Abcam) according to the manufacturer’s instructions. Chemiluminescence was imaged using a BioSpectrum 810 imaging system (UVP) with 3 min exposure. Location of the 40 cytokines detected by the array membranes are shown in Supplementary Table 2.

Densitometry of the spots in the array membranes was performed using ImageJ software (NIH). Briefly, the mean gray value of each spot was measured from all membranes using the same circular region of interest. The spots of interests were then subtracted from background (average of all negative control spots) and normalized to the positive control spots of the reference membrane (control FLS media). A fold change value was obtained by dividing normalized intensities of the membrane of interest and the control membrane, analyte-by-analyte.

### Culture of human FLS

FLS was obtained from 1 OA and 3 RA female patients (age range 61 – 81 years, see Supplementary Table 3 for detailed patient information) with approval from local ethics committee (07/H0809/35). These were then seeded separately at 10,000/well in 200 µl of DMEM media (Sigma) supplemented with 10% FCS (Fisher Scientific), 100 units/ml of Penicillin/Streptomycin (Fisher Scientific), 2% glutamine (Thermo Fisher) and fungizone (Thermo Fisher) into a 96-well flat bottom plate (Starlab). The plate was incubated overnight at 37 °C to allow for SF attachment and the following morning media was replaced with either 200µl of fresh DMEM containing 10 ng/ml of TNF-α (PeproTech), or fresh DMEM alone. Supernatants were collected following 24-hours of incubation at 37 °C.

### RNA extraction and RT-qPCR of human FLS

RNA was extracted using a RNeasy Micro Kit (Qiagen) according to the manufacturer’s protocol, then reverse transcribed and amplified using a slightly modified version of the Smart-seq2 method [48]. Briefly, in a RNase/DNA-free fume hood 2 µl of RNA was mixed with 1 µl of dNTP mix (Thermo Scientific), 1 µl of Oligo-dT primer (Merck) and 5.7 µl of RT mix, then reverse transcribed on a PTC-225 Gradient Thermal Cycler (MJ Research). 15 µl of PCR mix was added to the resulting cDNA and amplified according to previously established protocol. Concentrations were checked on a Qubit 3.0 (Invitrogen), with values ranging from 20.4 – 45.0 ng/µl. Samples were diluted to 1 ng/µl with double distilled H_2_O (ddH_2_O) and 1 ng was used for standard SYBR Green (Roche) RT-qPCR reactions on a LightCycler480 (Roche) to probe for the genes of interest (see Supplementary Table 1 for primer sequences). All primers were tested for their efficiency and specificity. The mean of housekeeping genes *B2M* and *Ywhaz* was used to calculate ΔCt values. All reactions were run in duplicate with water used as a negative control.

### DRG neuron isolation and culture

Lumbar DRG (L2-L5, those that primarily innervate the knee) were collected from FB labelled mice 7-10 days after knee injections in ice cold dissociation media containing L-15 Medium (1X) + GlutaMAX-l (Life Technologies), supplemented with 24 mM NaHCO_3_. DRG were then enzymatically digested in type 1A collagenase (Sigma) and trypsin solution (Sigma) at 37 °C, followed by mechanical trituration as described before [13]. Dissociated DRG neurons were plated onto poly-D-lysine and laminin coated glass bottomed dishes (MatTek, P35GC-1.5-14-C) and cultured either on their own (mono-culture), on a layer of FLS (co-culture, see below) or in 48-hour conditioned media from TNF-FLS for 24-hours. The DRG culture medium contained L-15 Medium (1X) + GlutaMAX-l, 10% (v/v) fetal bovine serum, 24 mM NaHCO_3_, 38 mM glucose, 2% (v/v) penicillin/streptomycin.

### DRG neuron/FLS co-culture

For co-culture studies, FLS were plated onto MatTek dishes and cultured for 24-hours with FLS culture media (with or without TNF-α stimulation). The next day medium was removed from FLS plates, then DRG neurons were isolated as described above and plated on top of the FLS. Co-culture plates were then kept in DRG culture medium for up to 24-hours for electrophysiological recording.

### Cell staining

#### FLS immunocytochemistry

FLS were plated overnight in wells of a 24-well plate, fixed with Zamboni’s fixative [57] for 10 min, permeabilized with 0.05 % TritonX-100 and blocked with antibody diluent (0.2% (v/v) Triton X-100, 5% (v/v) donkey serum and 1% (v/v) bovine serum albumin in PBS) for 30 min. The cells were then incubated overnight at 4 °C in 1:100 (in antibody diluent) anti-cadherin-11 antibody (CDH-11, rabbit polyclonal, Thermo Fisher, 71-7600). Cells were washed three times with PBS-tween and incubated in the conjugated secondary antibody, anti-rabbit Alexa-568 (1:1000 in PBS, Thermo Fisher, A10042) for 1-hour at room temperature (21 °C). The secondary antibody was washed off three times with PBS-tween and the cells were incubated in the nuclear dye DAPI (1:1000 in PBS, Sigma, D9452) for 10 min. Cells were further washed with PBS-tween once and imaged in PBS using an EVOS FLoid Cell Imaging Station (Thermo Fisher) at 598 nm (for CDH-11) and 350 nm (for DAPI) wavelength of light. Cells without primary antibody did not show fluorescence.

#### DRG neuron/FLS co-culture live cell stain

To visualize DRG neuron/FLS co-culture, live cell imaging was performed. FLS were plated on MatTek dishes with 1:1000 (diluted in FLS culture medium) CellTracker Deep Red Dye (Thermo Fisher, C34565) and incubated for 24-hours in a humidified 37 °C, 5% CO_2_ incubator. Dissociated DRG neurons (see above) were incubated in CellTracker Green Dye (1: 1000 diluted in DRG culture media, Thermo Fisher, C7025) for 15 min at room temperature (21 °C), centrifuged (16000 g, 3 min, 5415R, Eppendorf) and re-suspended in fresh medium. The FLS dishes were then washed twice with PBS and the neuronal suspension was plated on top of the FLS monolayer and incubated overnight in the incubator. The co-culture dishes were washed with PBS and imaged the following day using an Olympus BX51 microscope and QImaging camera at 650 nm (for deep red dye) and 488 nm (for green dye) wavelength of light.

### Whole-cell patch-clamp electrophysiology

DRG neurons were bathed in extracellular solution containing (ECS, in mM): NaCl (140), KCl (4), MgCl_2_ (1), CaCl_2_ (2), glucose (4) and HEPES (10) adjusted to pH 7.4 with NaOH. Only FB labelled neurons identified by their fluorescence upon excitation with a 365 nm LED (Cairn Research) were recorded. Patch pipettes of 5–10 MΩ were pulled with a P-97 Flaming/Brown puller (Sutter Instruments) from borosilicate glass capillaries and the intracellular solution used contained (in mM): KCl (110), NaCl (10), MgCl_2_ (1), EGTA (1), HEPES (10), Na_2_ATP (2), Na_2_GTP (0.5) adjusted to pH 7.3 with KOH.

Action potentials (AP) were recorded in current clamp mode without current injection for 20 s (to investigate spontaneous AP firing) or after step-wise injection of 80 ms current pulses from 0 – 1050 pA in 50 pA steps using a HEKA EPC-10 amplifier (Lambrecht) and the corresponding Patchmaster software. AP properties were analyzed using Fitmaster software (HEKA) or IgorPro software (Wavemetrics) as described before [13] and shown in Figure 3D (inset). Neurons were excluded from analysis if they did not fire an AP in response to current injection. For recording whole-cell voltage-clamp currents in response to TRP agonists capsaicin (1 μM from a 1 mM stock in ethanol, Sigma-Aldrich), cinnamaldehyde (100 μM from a 1 M stock in ethanol, Alfa Aesar) and menthol (100 μM from a 1 M stock in ethanol, Alfa Aesar), a 5 s baseline was established with ECS, followed by a 5 s randomized drug application. Peak drug response was measured in Fitmaster by subtracting the average of 2 s baseline immediately preceding the drug application and the maximum peak response reached during the 5 s of drug application. Peak current density is represented in graphs by dividing this peak response by the capacitance of the neuron. Data from at least four mice were used in all conditions (with each mouse being used for at least two conditions) and at least three neurons were recorded from each mouse in each category.

**Figure 1:**
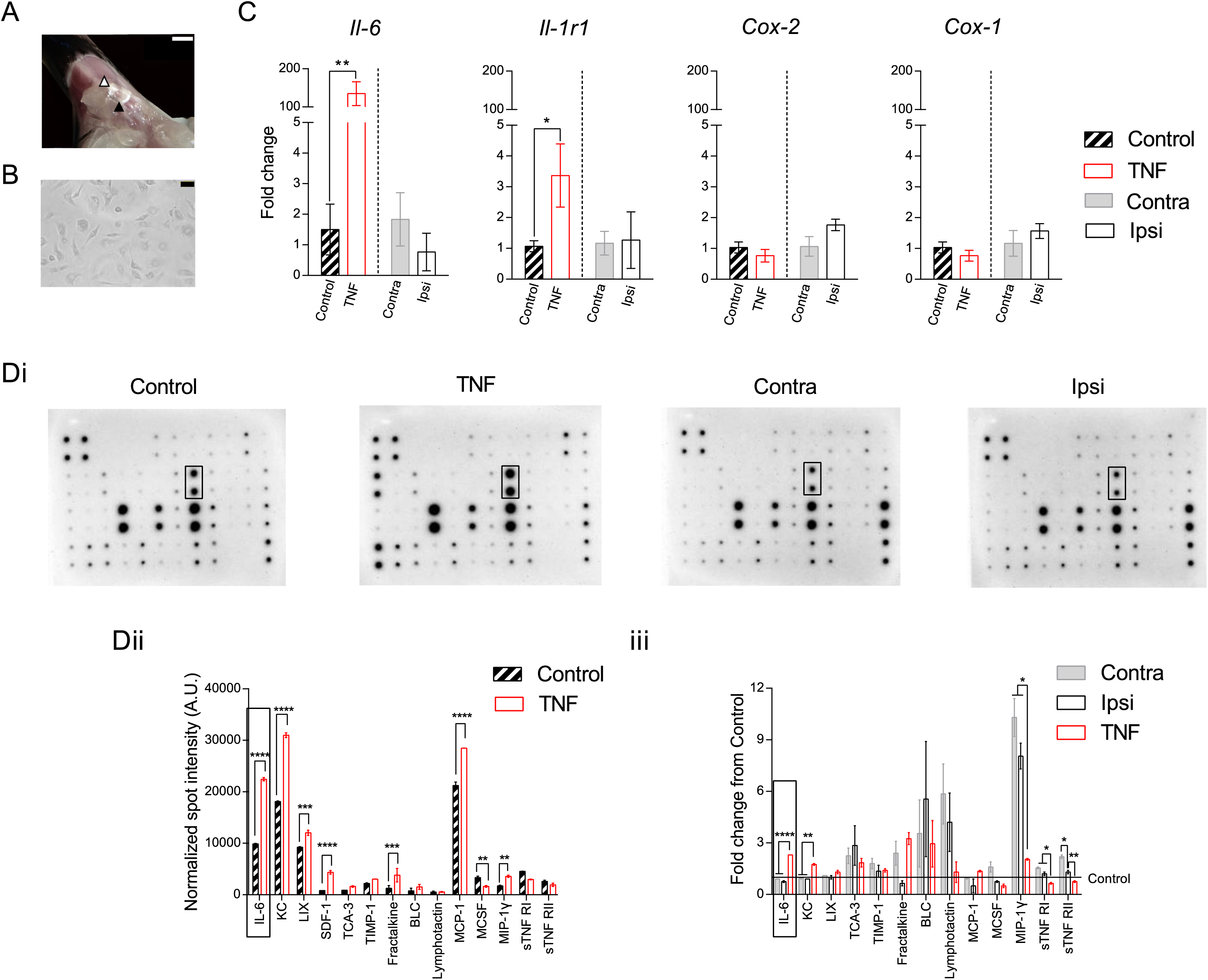
Induction of inflammation in FLS derived from mouse knee. A) Representative picture showing exposed inside of the patella (black triangle) and the surrounding ligament and fat pad (white triangle) after midline resection and distal pull of the quadriceps muscles. Scale bar = 2 mm. B) Representative picture of FLS in culture. Scale bar = 50 µm. C) Bars represent fold change of the genes *Il-6, Il-1r1, Cox-2* and *Cox-1* from either Contra (vs. Ipsi) or control (vs. TNF). Black hatched bars = control FLS, red bars = TNF-FLS, grey bar = Contra, white bar = Ipsi. All FLS at P5. D) Images of mouse inflammatory array membranes probed against FLS conditioned medium (i) which were quantified by densitometry and represented as bar graphs showing fold change of various cytokines between control and TNF-FLS (ii, multiple t-test with Holm-Sidak correction) and among Contra, Ipsi and TNF from control FLS media (iii, ANOVA with Tukey’s post-hoc test) and spot intensity differences. Dotted rectangles highlight IL-6 spots (Di) and corresponding quantifications (Dii, iii). Only cytokines that were present in all of the compared groups are shown in graphs. * p < 0.05, ** p < 0.01, *** p < 0.001 and **** p < 0.0001. Error bars = SEM.

**Figure 2.**
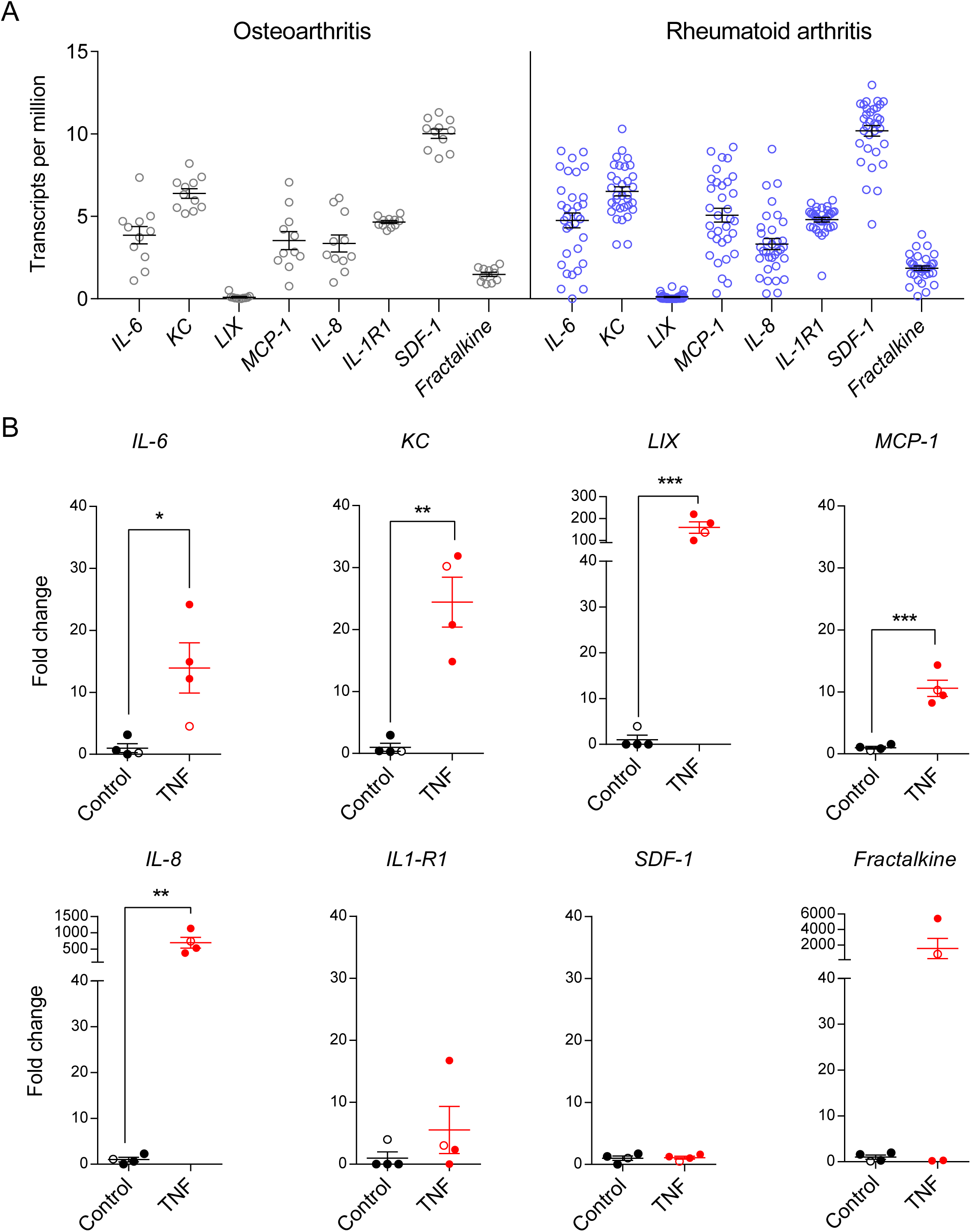
Pro-inflammatory gene expression in FLS derived from arthritic human patients. A) Transcripts per million values of selected inflammatory markers extracted from RNA-Seq data of human FLS from RA and OA donors [68]. Results reveal detectable levels of the pro-inflammatory genes *IL-6, IL-8, IL-1R1, KC, MCP-1, SDF-1* and *Fractalkine* in FLS from both OA (black) and RA (blue) patients. *LIX* was considered undetectable as the mean transcript per million value was below the threshold of 1. B) Bar graphs showing fold change in expression levels (determined by qPCR) of *IL-6, KC, LIX, MCP-1, IL-8, IL-1R1, SDF-1* and *Fractalkine* in control vs. TNF-α stimulated FLS derived from human RA (filled circles) and OA (open circles) patients (n = 4). * p <0.05, ** p <0.01, unpaired t-test.

**Figure 3:**
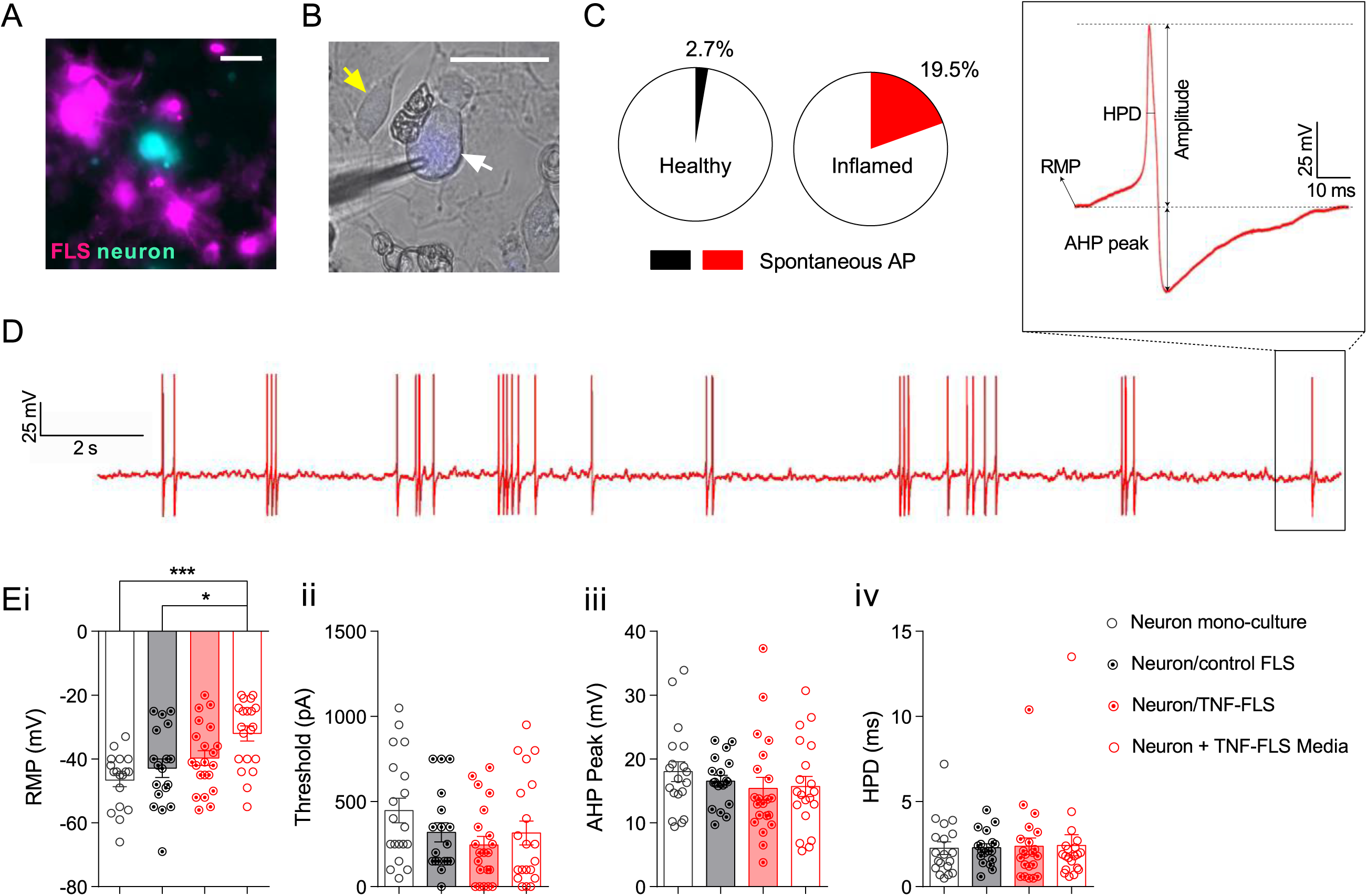
Murine TNF-FLS mediated increase in excitability of knee neurons. A) Representative live-cell imaging picture showing FLS (magenta) and neuron (cyan) in co-culture. Scale bar = 50 µm. B) Representative FB labelled knee neuron (white arrow) being recorded using a patch pipette (triangular shadow), surrounded by FLS (yellow arrow). Scale bar = 50 µm. C) Pie-chart showing proportion of knee neurons that fired AP without current stimulation in healthy (neuron mono-culture + neuron/control FLS, n = 37, black) and inflamed (neuron/TNF-FLS + neuron/TNF-FLS media, n = 41, red) condition. D) Representative knee neuron incubated with TNF-FLS media firing spontaneous AP along with schematic diagram of AP properties measured (inset). E) Bar graphs showing measured RMP (i), threshold (ii), AHP peak (iii) and HPD (iv) from knee neurons in mono-culture (n = 19, white bar/black open circle), in co-culture with control FLS (n = 18, grey bar/black dotted circle), in co-culture with TNF-FLS (n = 20, light red bar/red dotted circle) and incubated in TNF-FLS media (n = 21, white bar/red open circle). * p < 0.05 and *** p < 0.001, ANOVA followed by Tukey’s post hoc test. Data from 4-5 female mice in each group. Error bars = SEM.

### Ca^2+^ imaging

FLS or DRG neurons were incubated with the Ca^2+^ indicator, Fluo-4 AM (10 μM diluted in ECS from a 10 mM stock solution in DMSO, Invitrogen) for 30 min at room temperature (21 °C). Culture dishes were then washed and imaged using an inverted Nikon Eclipse T*i* microscope. Fluo-4 fluorescence was excited using a 470 nm LED (Cairn Research) and captured with a camera (Zyla cSMOS, Andor) at 1 Hz with 50 ms (for neurons) 250 ms (for FLS) exposure time using Micro-Manager software (v1.4; NIH). Solutions were perfused in this system through a gravity-driven 12 barrel perfusion system [20]. During imaging of neurons, MIP-1γ (10 and 100 ng/ml diluted in ECS from a stock concentration of 0.1 mg/ml in 0.1% bovine serum albumin/sterile PBS, Sigma) was applied for 20 s after establishing a baseline with ECS for 10 s. Neurons were allowed to recover for 5 minutes between drug applications. 50 mM KCl was used as a positive control.

During imaging of FLS, a 10 s baseline was established with ECS and then pH 4-7, and TRP agonists, 1 μM capsaicin, 100 μM cinnamaldehyde and 100 μM menthol were applied for 10 s before a wash out period of 4 min between drug applications. TRP agonist sensitivity and acid sensitivity was assessed on separate dishes. Three biological repeats were conducted on separate days for each condition. 10 μM ionomycin was used as a positive control after all FLS Ca^2+^ imaging experiments.

Analysis was conducted as described before [12]. Briefly, mean grey values were extracted by manually drawing around FLS or neurons in the ImageJ software. These values were then fed into a custom-made R-toolbox (https://github.com/amapruns/Calcium-Imaging-Analysis-with-R.git) to compute the proportion of cells responding to each drug and their corresponding magnitude of response (normalized to their peak ionomycin (FLS) or KCl (neuron) response, (ΔF/Fmax); cells not crossing threshold for positive controls were excluded from the analysis).

### Extraction of RNA-Seq data from RA and OA patient synovial tissue samples

Publicly available single cell RNA-Seq data of synovial tissue from 51 OA and RA patients [68] was used to investigate the presence of pro-inflammatory genes. For each sample, we extracted the transcripts per million (TPM) values for the same genes investigated for RT-qPCR of human FLS and organised data by patient ID, cell type and patient group. The 17 samples that failed or were pending quality control were excluded. The remaining 151 samples were subsequently plotted by dividing into two groups – OA and RA.

### Statistics

Comparisons between two groups with distributed variables were performed using two-sided Student’s t-tests (paired if comparing two conditions of the same sample, unpaired otherwise) with suitable corrections and amongst three groups using one-way analysis of variance (ANOVA) followed by Tukey’s post-hoc tests. Proportions were compared for categorical data using chi-sq tests. Data are shown as mean ± SEM.

## Results

### 1. TNF-FLS, but not FLS derived from CFA-induced inflamed knees of mice, have a pro-inflammatory phenotype and secrete cytokines that activate DRG neurons

Key genes have been described that act as identifiers of cultured FLS upon isolation by enzymatic digestion of mouse joints [29], as well as for establishing their inflammatory phenotype. In this study, we performed RT-qPCR on adherent cells (passages P2-P5) originating from mouse patellae that proliferated in culture (Figure 1A, B). From P2 to P3, the expression of the macrophage marker [5,29] cluster of differentiation 68 (*Cd*68, relative to the macrophage cell line RAW 267.4) was significantly reduced, whereas the expression of FLS markers *Cdh11* and *Cd248* [5,29] remained consistent from P2 to P5 (Supplementary Figure 1A, B showing expression of FLS marker genes); the endothelial marker *Cd31* was not detected (data not shown). These results suggest that from P3, cells cultured from mouse patellae were predominantly FLS and hence subsequent studies were conducted on FLS from P3-P6.

Concurrently, to establish the pro-inflammatory phenotype of FLS, we used RT-qPCR to determine the expression of the inflammatory genes *Il-6, Il-1r1* and *Cox-2*, as well as the constitutively expressed gene *Cox-1.* When control FLS (P5, n = 3 biological replicates) were stimulated with 10 ng/ml TNF-α for 48-hours, there was an upregulation of *Il-6* (fold changes: Control, 1.5 ± 0.8 vs. TNF-FLS, 135.1 ± 31.1, p = 0.006, unpaired t-test) and *Il-1r1* (fold changes: Control, 1.1 ± 0.2 vs. TNF-FLS, 3.4 ± 1.0, p = 0.04, unpaired t-test) expression levels, but not that of *Cox-2* (fold changes: Control, 1.0 ± 0.2 vs. TNF-FLS, 0.8 ± 0.2, p = 0.2, unpaired t-test) or *Cox-1* (fold changes: Control, 1.0 ± 0.2 vs. TNF-FLS, 0.8 ± 0.2, p = 0.18, unpaired t-test) (Figure 1C). However, when FLS derived from the inflamed knee (Ipsi, knee width, pre-CFA, 3.1 ± 0.09 mm vs. post-CFA, 4.1 ± 0.08 mm, n = 6, p = 0.0001, paired t-test, Supplementary Figure 1C showing knee width of mice pre and post-CFA injection) of CFA injected mice were compared to those of the matched contralateral knee (Contra), we did not find any changes in expression levels of the genes between Contra and Ipsi FLS across P2-P5 (Figure 1C, Supplementary Figure 1C).

We next tested the media (n = 3, each) isolated from control, Contra, Ipsi and TNF-FLS against a mouse inflammation antibody array (Figure 1D) to determine the levels of different secreted pro-inflammatory mediators. This demonstrated that when compared to control FLS media, TNF-FLS media showed presence of regulated on activation, normal T-cell expressed and secreted (RANTES) and granulocyte-macrophage colony stimulating factor (GM-CSF). Additionally, TNF-FLS contained higher levels of IL-6 (p < 0.0001), keratinocyte chemoattractant (KC, mouse homolog of IL-8, p < 0.0001), lipopolysaccharide induced chemokine (LIX, p < 0.0001), stromal cell derived factor 1 (SDF-1, p < 0.0001), fractalkine (p < 0.0001), monocyte chemoattractant protein 1 (MCP-1, p < 0.0001) and macrophage inhibitory protein 1γ (MIP-1γ, p = 0.002) and lower levels of macrophage colony stimulating factor (MCSF, p = 0.003) compared to control FLS media (multiple t-tests with Holm-Sidak corrections, Figure 1Dii). The TNF-α concentration in TNF-FLS media 48-hours after 10 ng/ml TNF-α stimulation was similar to the control media as determined by the inflammatory antibody array. Mirroring the absence of pro-inflammatory gene upregulation in Ipsi vs. Contra FLS, the spot intensity values of inflammatory cytokines between Ipsi and Contra FLS were similar (multiple t-tests with Holm-Sidak corrections, graph not shown), suggesting similar levels of secreted cytokines. Finally, when spot intensities were normalized to control FLS, TNF-FLS media showed increased levels of secreted IL-6 (p < 0.0001, ANOVA followed by Tukey’s multiple comparison test) and KC (p = 0.001, ANOVA followed by Tukey’s multiple comparison test) compared to Ipsi and Contra FLS (Figure 1Diii).

MIP-1γ was elevated in both Ipsi and Contra FLS when compared to control and TNF-FLS (Figure 1Diii), perhaps indicating its role in acute, systemic inflammatory pathways. MIP-1γ has been described as having a role in hyperalgesia induced by diabetic neuropathy [49] and OA-related pain [19]. We find that MIP-1γ directly activates mouse DRG neurons by producing a dose-dependent Ca^2+^ influx (Supplementary Figure 1D showing intracellular Ca^2+^ mobilization in DRG neurons upon MIP-1γ application), which thus might be a potential mechanism for the *in vivo* effects described above. However, because MIP-1γ does not have a human homolog (NCBI gene database entry #20308), and thus has limited clinical potential, we did not explore this further.

Taken together, our data suggest that TNF-α stimulation of FLS upregulates expression of canonical inflammatory markers, whereas the knee inflammation observed 24-hours after intra-articular CFA injection does not correlate with a sustained pro-inflammatory phenotype of FLS isolated from patellae of CFA mice. We also identify other soluble mediators secreted by TNF-FLS that might be involved in FLS-DRG neuron communication. Therefore, given the pro-inflammatory phenotype of TNF-FLS (and the lack of such phenotype in the FLS expanded from CFA-injected knees), we focused solely on the TNF-FLS for the rest of the study.

### 2. TNF-α stimulated human FLS from OA and RA patients show increased expression of multiple pro-inflammatory genes that were upregulated in mouse TNF-FLS

In order to understand whether the identified pro-inflammatory mediators from mouse FLS are important in human arthritis, we examined gene expression in human FLS. In arthritis, activated FLS contribute to pathogenesis by damaging synovial membranes and secreting inflammatory cytokines which in turn recruit immune cells [32,44,45]. In line with these reports, data re-plotted from a recently published single-cell RNA-seq dataset of FLS from 51 OA and RA patients [68] confirms detectable levels of pro-inflammatory genes *IL-6, KC, MCP-1, IL-8, IL1-R1, SDF-1* and *Fractalkine* (Figure 2A), which matches our mouse data in Figure 1.

Furthermore, to establish whether a similar pro-inflammatory phenotype can be observed in TNF-α treated human FLS as in mouse FLS, we used RT-qPCR. When four batches of human FLS derived from four separate patients were stimulated with 10 ng/ml TNF-α for 24-hours, there was increased expression of *IL-6* (p = 0.02), *KC* (p = 0.001), *LIX* (p = 0.009), *MCP-1* (p = 0.0004) and *IL-8* (p = 0.005) compared to unstimulated controls (unpaired t-test) (Figure 2B). However, expression of *SDF-1* (p = 0.8), *IL-1R1* (p = 0.3) and *Fractalkine* (p = 0.3) was not significantly different in the TNF group compared to control (unpaired t-test), although for the latter two genes, FLS from some patients showed considerable increase after TNF-α stimulation (Figure 2B).

The data presented here demonstrate the translational potential of our data derived from mouse FLS (Figure 1), as well as supporting a previous study that showed increased cytokine production from TNF-α stimulated human FLS from both normal and RA donors compared to unstimulated ones [32].

### 3. TNF-FLS derived from mice increase knee-innervating DRG neuron excitability in co-culture

Changes in primary sensory neuron excitability underlie peripheral sensitization which drives arthritis-related pain [58]. Non-neuronal cells like FLS, which are in close proximity with distal terminals of knee neurons, can play instrumental roles in modulating sensory neuron excitability by direct cell contact and/or secretion of pro-inflammatory cytokines. To determine neuron/FLS communication in health and inflammation we compared the DRG neuronal excitability of four groups: 1) knee neuron mono-culture, 2) knee neurons co-cultured with control FLS (Figure 3A, B), 3) knee neurons with FLS that have been exposed to TNF media for 24-hour followed by DRG culture media for 24-hours and 4) knee neurons with conditioned media from FLS that have been exposed to TNF media for 48-hours to understand the role of soluble mediators in peripheral sensitization. In current-clamp mode, we observed many neurons spontaneously firing AP during a 20 s recording period (mostly with an intermittent firing pattern, although neurons that fired >100 APs had a continuous firing pattern) in groups 3 and 4. In order to statistically compare the proportions, we converted our four group data into binary categories: neuron mono-culture and neuron/control FLS co-culture were assigned to the class “healthy”, and neuron/TNF-FLS and neuron/TNF-FLS media were assigned to the class “inflamed”. 19.5% of inflamed neurons (neuron/TNF-FLS, 5/21; neuron/TNF-FLS media, 3/19) fired spontaneous AP compared to 2.7% (neuron mono-culture, 0/19; neuron/control FLS, 1/18) of healthy neurons (p = 0.02, chi-sq test). The mean number of APs fired (within the 20 s recorded without current injection) by neurons in the neuron/TNF-FLS and neuron/TNF-FLS media categories was 31.6 ± 18.6 and 76.0 ± 62.5 respectively, while the neuron in the neuron/control FLS group fired 18 times. This result suggests that there is a general increase in excitability of knee neurons when exposed to an FLS-mediated inflammatory environment (Figure 3C, D).

Upon measuring the AP properties (Figure 3D inset) we found that the resting membrane potential (RMP) was more depolarized in the neuron/TNF-FLS media group compared to both neuronal mono-culture (p = 0.0004) and neuron/control FLS co-culture (p = 0.012), which highlights that secreted pro-inflammatory factors from TNF-FLS likely act upon knee neurons to increase their excitability (ANOVA followed by Tukey’s post hoc comparison, Figure 3Ei). This depolarizing trend in RMP was also observed when the data were divided into small and medium-large neurons (Supplementary Figure 2). The other measured properties were unchanged among the four groups (ANOVA followed by Tukey’s post hoc comparison, Table 1, Figure 3Eii-iv).

**Table 1:** AP properties of knee neurons. *** represents p < 0.001 when compared to knee neuron mono-culture, $ represents p < 0.05 when compared to neuron/control FLS co-culture.

|  | Knee neuron mono-culture<br>(n = 19) |  | Knee neuron/control FLS<br>(n = 18) |  | Knee neuron/TNF-FLS<br>(n = 21) |  | Knee neuron/TNF-FLS media<br>(n = 20) |  |
| --- | --- | --- | --- | --- | --- | --- | --- | --- |
|  | Mean | SEM | Mean | SEM | Mean | SEM | Mean | SEM. |
| RMP (mV) | -46.6 | 2.0 | -42.8 | 2.9 | -39.7 | 2.3 | -32.0***/\$ | 2.3 |
| Threshold (pA) | 447.4 | 71.5 | 319.4 | 56.3 | 245.2 | 49.4 | 315.0 | 70.0 |
| Half peak duration (HPD, ms) | 2.2 | 0.4 | 2.3 | 0.2 | 2.4 | 0.5 | 2.4 | 0.6 |
| Afterhyperpolarization amplitude (AHP peak, mV) | 13.7 | 2.6 | 19.6 | 2.7 | 16.1 | 2.9 | 21.6 | 3.3 |

### 4. TNF-FLS derived from mice increase TRPV1 function and decrease TRPA1/TRPM8 function of DRG neurons in co-culture

DRG neurons in co-culture with inflamed FLS derived from AIA rats reportedly have increased expression of TRPV1 [4]. Here we sought to investigate whether FLS can modulate knee neuron responses to TRP channel agonists using whole-cell patch-clamp recordings. Knee neurons in mono-culture and when co-cultured with control FLS displayed very similar responses, i.e. a mean capsaicin peak current density response of 3.5 ± 1.0 pA/pF (n = 7/19) and 3.2 ± 0.9 pA/pF (n = 7/18) respectively. By contrast, knee neurons in neuron/TNF-FLS co-culture showed a trend for larger magnitude of capsaicin responses (27.7 ± 9.3 pA/pF, n = 8/21), and those cultured with TNF-FLS media had a capsaicin response of 65.8 ± 25.7 pA/pF (n = 7/20) which was significantly larger in magnitude than neuronal mono-culture (p = 0.02) and neuron/control FLS co-culture (p = 0.02) (ANOVA followed by Tukey’s post-hoc comparison, Figure 4Ai,ii). However, no difference was observed between the percentage of capsaicin responders in healthy (40.5%) vs. inflamed groups (37.5%, p = 0.8, chi-sq test, Figure 4Aiii), suggesting that although TRPV1 channel function is sensitized in TRPV1-expressing neurons, TNF-FLS exposure does not induce TRPV1 expression in neurons lacking any basal expression.

**Figure 4:**
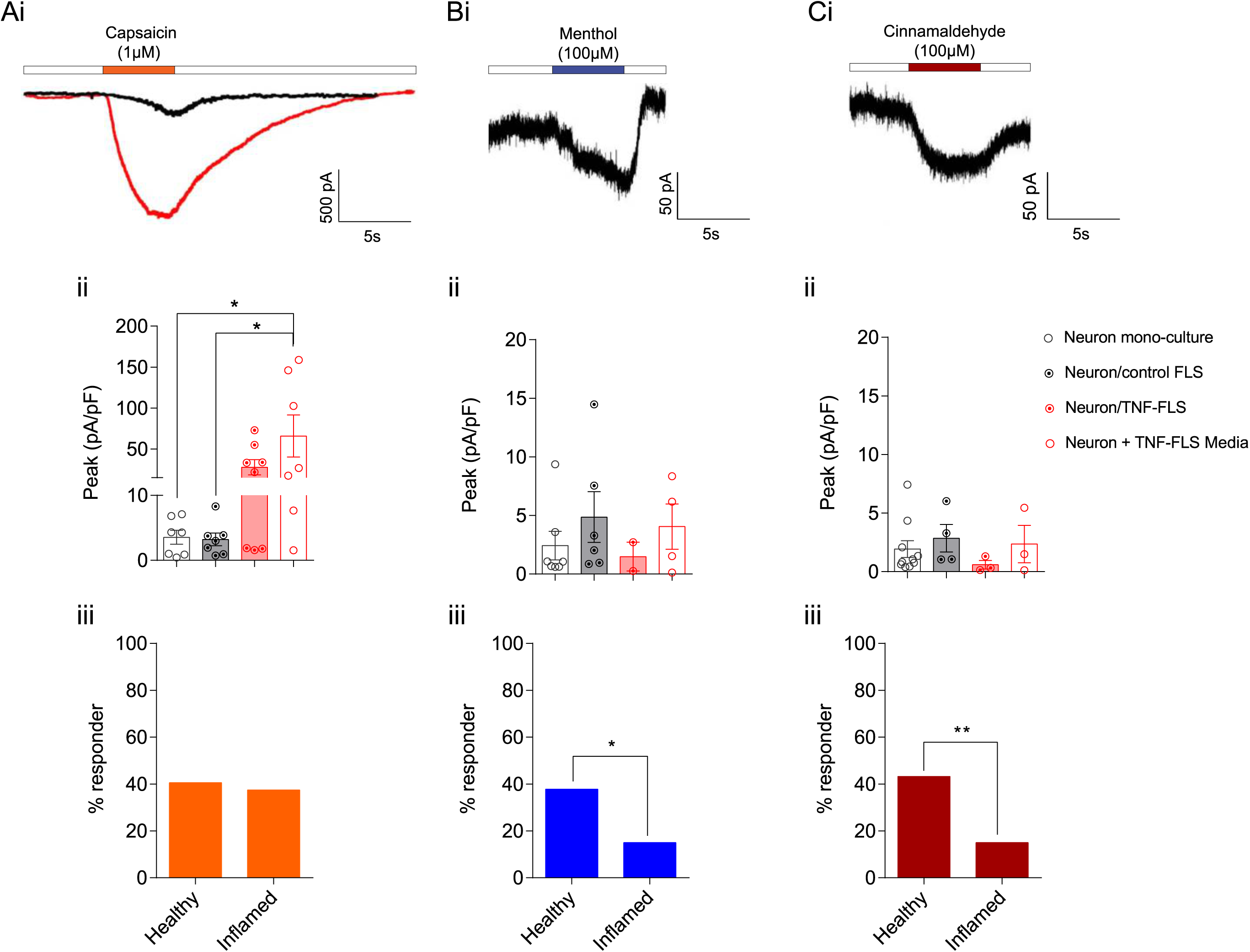
Murine TNF-FLS mediated modulation of TRP agonist response in knee neurons. Representative traces showing capsaicin (TRPV1 agonist, Ai), cinnamaldehyde (TRPA1 agonist, Bi) and menthol-(TRPM8 agonist, Ci) evoked responses in knee neurons. Black traces obtained from knee neuron in mono-culture, red trace obtained from knee neuron incubated in TNF-FLS media. White boxes represent perfusion of extracellular solution. Bar graphs showing peak current densities of capsaicin-(Aii), cinnamaldehyde-(Bii) or menthol-evoked (Cii) currents from knee neurons in mono-culture (white bar/black open circle), in co-culture with control FLS (grey bar/black dotted circle), in co-culture with TNF-FLS (light red bar/red dotted circle) and incubated in TNF-FLS media (white bar/red open circle) Comparison between groups made using ANOVA with Tukey’s post hoc test. Bar graphs showing percent of knee neurons that responded to capsaicin (Aiii), cinnamaldehyde (Biii) and menthol (Ciii) in healthy (neuron mono-culture + neuron/control FLS) and inflamed (neuron/TNF-FLS + neuron/TNF-FLS media) condition. Comparison made using chi-sq test. * p < 0.05. Data from 4-5 female mice in each group. Error bars = SEM.

With regards to menthol (mono-culture, 2.4 ± 1.2 pA/pF, n = 7/19; neuron/healthy FLS, 4.8 ± 2.1 pA/pF, n = 6/18; neuron/TNF-FLS, 1.4 ± 1.2 pA/pF, n = 2/21; neuron/TNF-FLS conditioned media, 4.0 ± 1.9 pA/pF, n = 4/20; Figure 4Bi) and cinnamaldehyde (mono-culture, 1.9 ± 0.7 pA/pF, n = 10/19; neuron/healthy FLS, 2.8 ± 1.1 pA/pF, n = 4/18; neuron/TNF-FLS, 0.6 ± 0.3 pA/pF, n = 3/21, neuron/TNF-FLS conditioned media, 2.3 ± 1.5 pA/pF, n = 3/20; Figure 4Ci) responses, all four groups showed similar mean peak current density values (Figure 4Bii, Cii). However, in response to both the TRPA1 (cinnamaldehyde) and TRPM8 (menthol) agonists, the percentage of responding neurons was significantly less in the inflamed group compared to healthy (menthol: healthy vs. inflamed, 37.8% vs. 15%, p = 0.02, chi-sq test; cinnamaldehyde: healthy vs. inflamed, 43.2% vs. 15%, p = 0.006, chi-sq test, Figure 4Biii, Ciii). Taken together our data suggest that “inflamed” FLS can alter TRP channel agonist response of knee neurons in co-culture.

Overall, our data suggest that pro-inflammatory mediators secreted from FLS are capable of increasing excitability and TRPV1 function of knee-innervating DRG neurons. It has been hypothesized that secretion by FLS is mediated by an increase in intracellular [Ca^2+^] [16,35] and depolarization. Furthermore, during inflammatory arthritis, FLS are located in an environment with abundant algogens that have been shown to signal through the modulation of TRP channels [9] and there is also mixed evidence in animals and humans for development of tissue acidosis [3,24,66]. TRP channels and acid sensors in FLS can increase intracellular [Ca^2+^] [26,35] and therefore, we tested the sensitivity of mouse knee-derived FLS to a range of acidic pH stimuli, as well as the prototypic TRP channel agonists, capsaicin (TRPV1), cinnamaldehyde (TRPA1) and menthol (TRPM8) using Ca^2+^ imaging. We found that both control and TNF-FLS responded to a range of pH solutions (pH 4.0 – 7.0) with an increase in intracellular [Ca^2+^], with an increased percentage of TNF-FLS responding to acid than control FLS at pH 5.0-6.0 (Supplementary figure 3). However, capsaicin (control, 3/201, TNF-FLS, 13/252), cinnamaldehyde (control, 1/201, TNF-FLS, 11/252) and menthol (control, 0/201, TNF-FLS, 1/194) evoked a Ca^2+^ response in only a few cells (data not shown), leading us to conclude that very few mouse FLS have functional TRPV1, TRPA1 and TRPM8 ion channels. RT-PCR for control and TNF-FLS revealed expression of multiple proton-sensors and low *Trpv1* expression, further validating the Ca^2+^ imaging results (Supplementary figure 3). Therefore, our data supports the hypothesis of Ca^2+^ dependent secretion in FLS leading to increased knee neuron sensitization and nociception.

## Discussion

FLS can respond to pro-inflammatory environments and then themselves become effectors for driving disease pathology, and hence are called “passive responders and imprinted aggressors” [9]. In support of their activated phenotype, here we demonstrate that FLS obtained from human OA and RA patients and from cell-outgrowth of mouse patella can respond to TNF-α stimulation to increase secretion and expression of several pro-inflammatory mediators. We used these findings to establish a co-culture system, which showed that murine FLS and knee neurons can interact, specifically, factors secreted by TNF-FLS increase knee neuron excitability and the magnitude of the response to capsaicin, whereas the proportions of cinnamaldehyde and menthol responding neurons are diminished. To the best of our knowledge, this is the first report to demonstrate FLS-mediated changes in neuronal excitability in co-culture and hence directly demonstrate how FLS can regulate articular neurons and in turn arthritis-related pain.

FLS from mouse are generally cultured by enzymatically digesting and combining excised joints of fore- and hind-limbs [26,29,50,56], which assumes similarity of FLS derived from all joints. However, a genome-wide study on DNA-methylation has shown that important differences exist between knee and hip FLS, including of genes involved in IL-6 signalling [1]. By using a cell-outgrowth method to culture mouse FLS, as previously described in humans [34] and rats [4], we have avoided the biological ambiguity introduced by joint-to-joint variability. Using these FLS, we investigated the expression of inflammatory genes *Il-6, Il-1r1* and *Cox-2*, all of which have been linked to the inflammatory phenotype of FLS. In brief: stimulating human-derived FLS with IL-1β increases *Cox-2* and *Il-6* expression [34]; increased *Il-6* expression is seen in FLS derived from K/BxN mice and following 24-hour stimulation of healthy mouse FLS with TNF-α [29]. Lastly, supernatants from cultured FLS derived from rats 3-days into AIA model show increased IL-6 and PGE_2_ [4]. Here we observed that neither the expression level of *Il-6, Il-1r1, Cox-2*, nor the level of secreted cytokines was upregulated in FLS derived from CFA-injected knee (our local ethics review body did not permit investigation of > 24-hours following CFA injection). This is possibly because the model used here is too brief to influence FLS gene expression, which is supported by a previous study where FLS derived from rats with longer (3-28 days) AIA-induced knee inflammation did have higher PGE_2_ and IL-6 concentrations in culture supernatants compared to control [4].

Therefore, we created a model of inflammation *in vitro* where we exposed FLS to TNF-α, a cytokine that is locally upregulated within 3-hours of intra-plantar CFA injection in mice [65]. It is also present in high concentrations in the synovial fluid [36,53] and tissue of OA and RA patients [47,61]; along with anti-TNF-α agents being a leading treatment of RA [60]. We show that TNF-α stimulated FLS display increased expression of *Il-6* and *Il-1r1* mRNA, with a concomitant increase in secretion of many pro-inflammatory cytokines, including IL-6 - a finding reported previously in human FLS [32] and replicated in this study. This observation is consistent with FLS being effectors of inflammatory arthritis, and hence we explored the functional repercussions of this pro-inflammatory phenotype by investigating the ability of “inflamed” FLS to induce functional changes in nerves supplying the knee joint to drive pain.

In order to understand the effector role of FLS in driving nociception through peripheral sensitization, we set up a co-culture system combining FLS and knee neurons. Studying co-culture of rat FLS and DRG neurons, von Banchet *et al* showed using immunohistochemistry that bradykinin 2 receptor labelling (but not that of neurokinin 1 or TRPV1) was increased when DRG neurons were cultured with healthy rat FLS [4], i.e. co-culture of DRG neurons with FLS can alter expression of genes associated with nociception. Therefore, we first verified that knee neurons in co-culture with control FLS do not show dysregulation of excitability or TRP agonist response. Then we asked whether TNF-FLS modulate knee neuron function and found using whole-cell patch clamp that 23% and 16% of knee neurons in neuron/TNF-FLS co-culture and neuron/TNF-FLS media (19.5 % in the combined “inflamed” condition) respectively evoked spontaneous AP compared to 6% in neuron/control FLS and 0% in neuron mono-culture (2.7% in the combined “healthy” condition). This suggests that TNF-FLS increase excitability of knee neurons and thus contribute to arthritic pain.

von Banchet *et al* [4] also showed that compared to mono-culture, FLS derived from acute and chronic AIA rats induced an increase in TRPV1 protein expression in DRG neurons, which can also lead to sensitization and hence arthritic pain. However, we did not observe an increase in the proportion of neurons responding to capsaicin between healthy and inflamed conditions, which might reflect a species difference and/or difference in knee-specific neuronal population (unlike in this study, von Banchet *et al* did not discriminate between knee-innervating neurons and non-knee innervating neurons); however, we did observe that capsaicin responsive neurons produced larger magnitude responses when incubated with TNF-FLS medium (see below). We also observed that cinnamaldehyde- and menthol-evoked responses were decreased in the inflamed condition, suggesting functional downregulation of TRPA1 and TRPM8, which might compensate for TRPV1 sensitization with regard to overall neuron excitability. Functional downregulation of TRPA1 might be explained through desensitization via increased TRPV1 function (see below) [2] or via an increase in intracellular [Ca^2+^] (due to increased excitability in the inflamed condition) [63]. The latter reason can also be applied to explain decrease of menthol-evoked responses [52].

We also observed a tendency of a more depolarized RMP and an enhanced magnitude of capsaicin-evoked peak current density of knee neurons in “inflamed” conditions, albeit both only reached statistical significance in the TNF-FLS media incubated group when compared to mono-culture and neuron/control FLS. However, we did not observe a decrease in the threshold of AP firing in the inflamed group, despite an increased proportion of these neurons firing spontaneously, thus suggesting perhaps differential susceptibility of certain knee neuron subgroups to TNF-FLS mediated sensitization. For example, a secreted factor may cause one neuronal subpopulation to fire spontaneously (i.e. an AP threshold of 0 pA), but overall the average AP threshold would not change due to other neuronal subgroups with a higher threshold not being susceptible to modulation by that secreted factor. Taken together, our data suggest soluble mediators released by FLS are key players in modulating knee neuron excitability, although cell contact with FLS might also play a role in sensitization. We were unable to compare the relative contributions of cell contact and soluble mediators to sensitization because, in this set-up TNF-FLS is always accompanied by TNF-FLS media. We posit that because our experimental design involved a media change 24-hours after TNF incubation of FLS (so as not to directly stimulate neurons with residual TNF-α [18]) in the neuron/TNF-FLS co-culture condition, the accumulated soluble mediator concentrations were lower compared to when knee neurons were incubated in 48-hours TNF-FLS conditioned media (no remaining TNF-α was measured in the media at this point). This decrease in concentration of TNF-α is hypothesized to be because of FLS’s inability to endogenously secrete TNF-α and/or due to protein degradation [38]. Our data is consistent with a previous study in FLS cells that measured TNF-α as <4 ng/ml 48-hours after stimulation with 10 ng/ml [38]. Similarly, in another non-TNF-α secreting cell line (Clara cells, from airways) 24-hours after initial stimulation with 20 ng/ml TNF-α, the remaining concentration of TNF-α fell below the detection level of the antibody array used [22].

Indeed, we have recently established the role of soluble mediators present in OA synovial fluid in increasing neuronal excitability and TRPV1 function [12]. These soluble mediators mainly consist of cytokines/chemokines which form complex signalling pathways with neurons (reviewed in [43,54]). In this study, we have identified several of them to be upregulated in TNF-FLS media which have been reported to be able to directly signal to neurons: IL-6 [23], KC [10], RANTES [46], GM-CSF [21], LIX [42], SDF-1 [46], MCP-1 [33], and MIP-1γ (present study). In the case of IL-6, MCP-1 and GM-CSF there have also been reports of an increase in TRPV1 function [21,23,33].

In summary, we have established a co-culture system to provide evidence for a direct inflammation-pain axis through FLS and DRG neurons. In the future, it can be adapted to investigate FLS-mediated peripheral sensitization using FLS obtained from chronic models of arthritis, activated by other pro-inflammatory cytokines or arthritic synovial fluid.

## Acknowledgements

The authors thank Dr Dora Lopresto (Department of Pharmacology, University of Cambridge) for technical help with cell staining and Drs Lucy Durham and Peter Earnshaw for collection and processing of human FLS samples.

## Conflict of interests

The authors declare no competing interests.

## Research Funding

This study was supported by Versus Arthritis Project Grants (RG 20930 and RG 21973), a Rosetrees grant (M818) to E. St. J. S. and G.C., Crohn’s and Colitis UK grant to D.C.B. (PC2019/1-Bulmer), and a King’s Together Award to L.S.T. and F.D. S.C. and C.N.B were supported by Gates Cambridge Trust scholarships. L.A.P. was supported by the University of Cambridge BBSRC Doctoral Training Programme (BB/M011194/1). S.L. was supported by Versus Arthritis (21139).

## Availability of data and materials

The datasets supporting the conclusions of this article will be available in University of Cambridge Apollo Repository (https://doi.org/10.17863/CAM.44367).

---

**Supplementary Figure 1.**
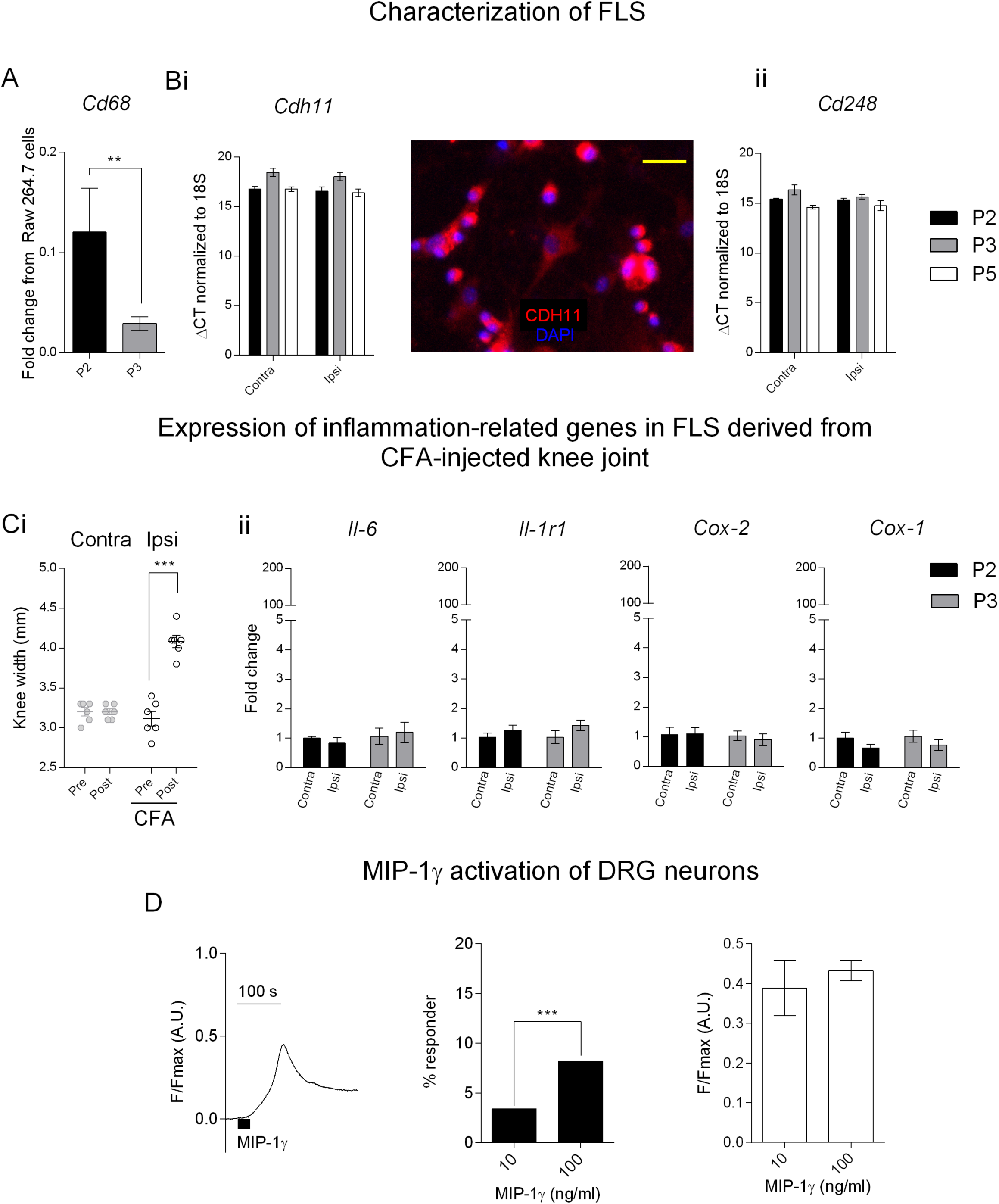

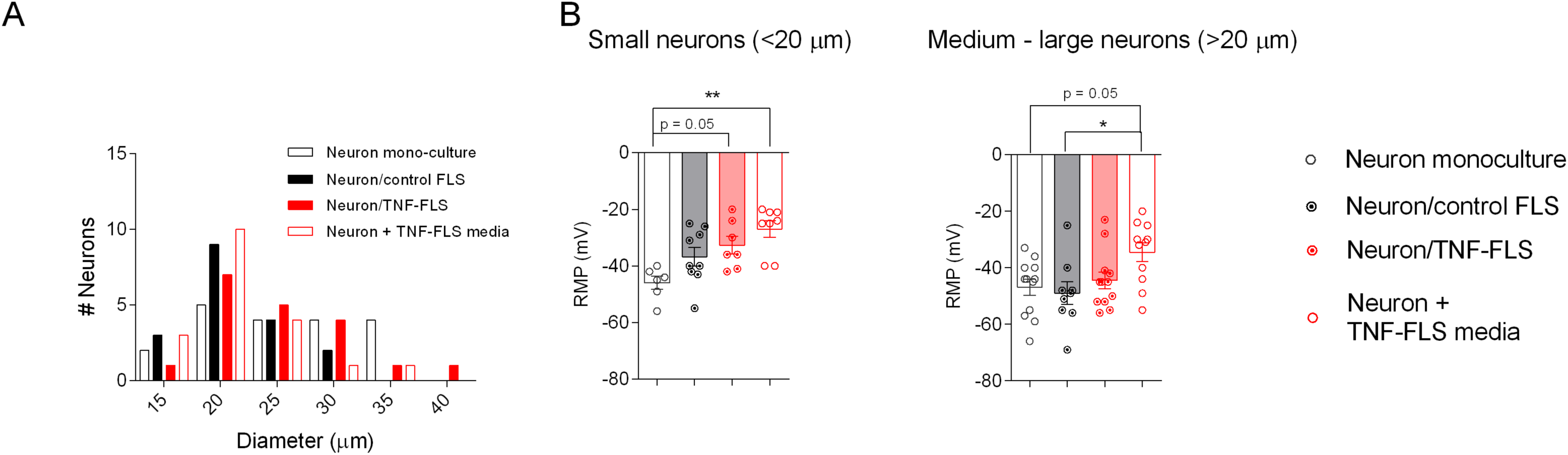

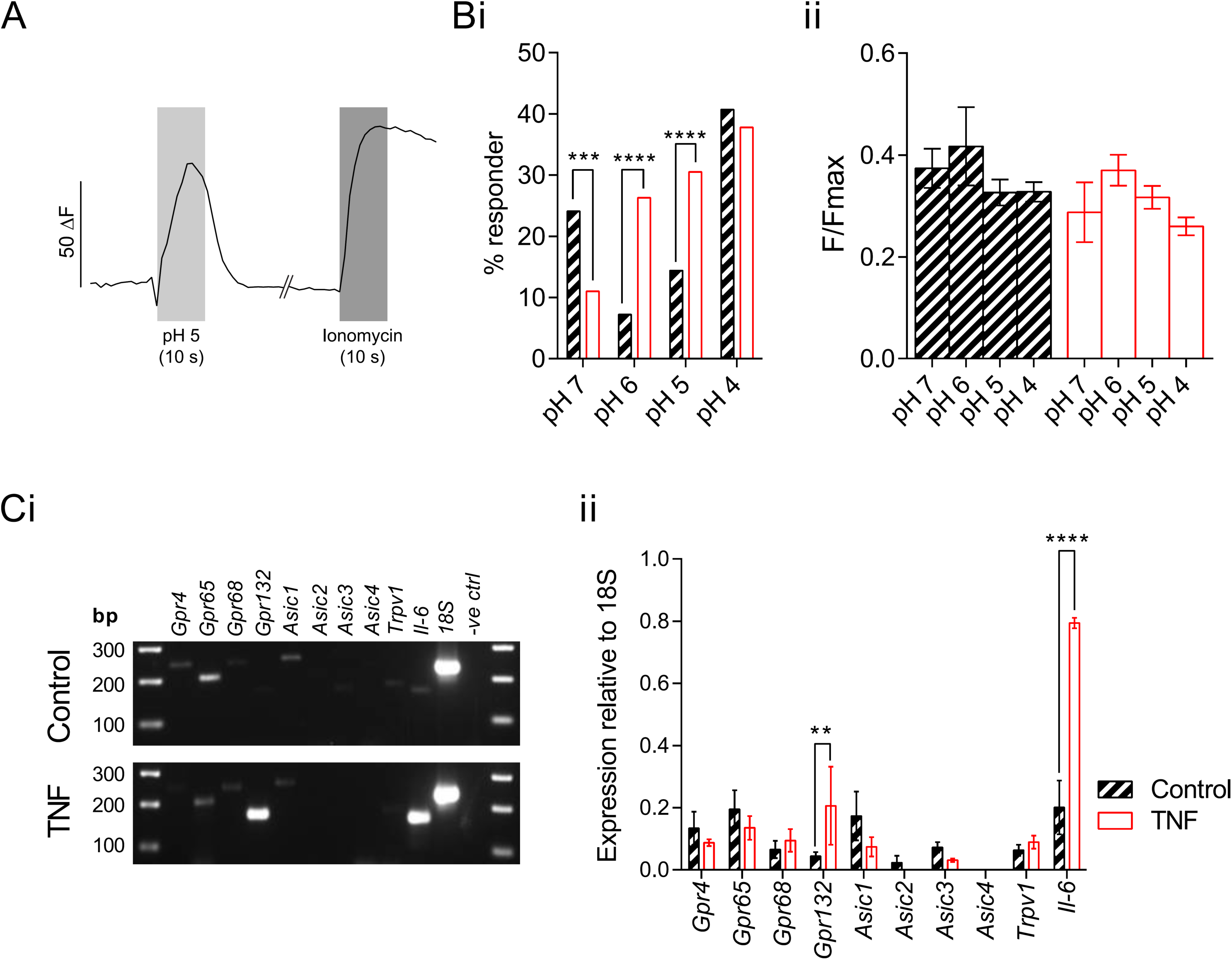

## Notes

#### Summary of Updates

Extra data and manuscript amendments following revision of article during peer review process.

## References

[1] Ai R, Hammaker D, Boyle DL, Morgan R, Walsh AM, Fan S, Firestein GS, Wang W. Joint-specific DNA methylation and transcriptome signatures in rheumatoid arthritis identify distinct pathogenic processes. Nat Commun 2016;7:11849–11849.

[2] Akopian AN, Ruparel NB, Jeske NA, Hargreaves KM. Transient receptor potential TRPA1 channel desensitization in sensory neurons is agonist dependent and regulated by TRPV1-directed internalization. J Physiol 2007;583:175–193.

[3] Andersson SE, Lexmuller K, Johansson A, Ekstrom GM. Tissue and intracellular pH in normal periarticular soft tissue and during different phases of antigen induced arthritis in the rat. J Rheumatol 1999;26:2018–2024.

[4] von Banchet GS, Richter J, Hückel M, Rose C, Bräuer R, Schaible H-G. Fibroblast-like synovial cells from normal and inflamed knee joints differently affect the expression of pain-related receptors in sensory neurones: a co-culture study. Arthritis Research & Therapy 2007;9:R6–R6.

[5] Bartok B, Firestein GS. Fibroblast-like synoviocytes: key effector cells in rheumatoid arthritis. Immunological reviews 2010;233:233–255.

[6] Blue ML, Conrad P, Webb DL, Sarr T, Macaro M. Interacting monocytes and synoviocytes induce adhesion molecules by a cytokine-regulated process. Lymphokine Cytokine Res 1993;12:213–218.

[7] Bombara MR, Webb DL, Conrad P, Marior CW, Sarr T, Ranges GE, Aune TM, Grave JM, Blue M-L. Cell contact between T cells and synovial fibroblasts causes induction of adhesion molecules and cytokines. Journal of Leukocyte Biology 1993;54:399–406.

[8] Bondeson J, Wainwright SD, Lauder S, Amos N, Hughes CE. The role of synovial macrophages and macrophage-produced cytokines in driving aggrecanases, matrix metalloproteinases, and other destructive and inflammatory responses in osteoarthritis. Arthritis Research & Therapy 2006;8:R187.

[9] Bottini N, Firestein GS. Duality of fibroblast-like synoviocytes in RA: passive responders and imprinted aggressors. Nat Rev Rheumatol 2013;9. doi: 10.1038/nrrheum.2012.190.

[10] Brandolini L, Benedetti E, Ruffini PA, Russo R, Cristiano L, Antonosante A, d’Angelo M, Castelli V, Giordano A, Allegretti M, Cimini A. CXCR1/2 pathways in paclitaxel-induced neuropathic pain. Oncotarget 2017;8:23188–23201.

[11] Burger D, Rezzonico R, Li J-M, Modoux C, Pierce RA, Welgus HG, Dayer J-M. Imbalance between interstitial collagenase and tissue inhibitor of metalloproteinases 1 in synoviocytes and fibroblasts upon direct contact with stimulated T lymphocytes: Involvement of membrane-associated cytokines. Arthritis & Rheumatism 1998;41:1748–1759.

[12] Chakrabarti S, Jadon DR, Bulmer DC, Smith EStJ. Human osteoarthritic synovial fluid increases excitability of mouse dorsal root ganglion sensory neurons: an in-vitro translational model to study arthritic pain. Rheumatology 2019. doi: 10.1093/rheumatology/kez331.

[13] Chakrabarti S, Pattison LA, Singhal K, Hockley JRF, Callejo G, Smith EStJ. Acute inflammation sensitizes knee-innervating sensory neurons and decreases mouse digging behavior in a TRPV1-dependent manner. Neuropharmacology 2018;143:49–62.

[14] Chen C-X, Chen J-Y, Kou J-Q, Xu Y-L, Wang S-Z, Zhu Q, Yang L, Qin Z-H. Suppression of Inflammation and Arthritis by Orally Administrated Cardiotoxin from Naja naja atra. Evid Based Complement Alternat Med 2015;2015:387094.

[15] Chomarat P, Rissoan MC, Pin JJ, Banchereau J, Miossec P. Contribution of IL-1, CD14, and CD13 in the increased IL-6 production induced by in vitro monocyte-synoviocyte interactions. J Immunol 1995;155:3645.

[16] Clark RB, Schmidt TA, Sachse FB, Boyle D, Firestein GS, Giles WR. Cellular electrophysiological principles that modulate secretion from synovial fibroblasts. J Physiol 2017;595:635–645.

[17] Croft AP, Campos J, Jansen K, Turner JD, Marshall J, Attar M, Savary L, Wehmeyer C, Naylor AJ, Kemble S, Begum J, Dürholz K, Perlman H, Barone F, McGettrick HM, Fearon DT, Wei K, Raychaudhuri S, Korsunsky I, Brenner MB, Coles M, Sansom SN, Filer A, Buckley CD. Distinct fibroblast subsets drive inflammation and damage in arthritis. Nature 2019;570:246–251.

[18] Czeschik JC, Hagenacker T, Schäfers M, Büsselberg D. TNF-α differentially modulates ion channels of nociceptive neurons. Neuroscience Letters 2008;434:293–298.

[19] Dawes JM, Kiesewetter H, Perkins JR, Bennett DLH, McMahon SB. Chemokine expression in peripheral tissues from the monosodium iodoacetate model of chronic joint pain. Mol Pain 2013;9:57–57.

[20] Dittert I, Benedikt J, Vyklický L, Zimmermann K, Reeh PW, Vlachová V. Improved superfusion technique for rapid cooling or heating of cultured cells under patch-clamp conditions. Journal of Neuroscience Methods 2006;151:178–185.

[21] Donatien P, Anand U, Yiangou Y, Sinisi M, Fox M, MacQuillan A, Quick T, Korchev YE, Anand P. Granulocyte-macrophage colony-stimulating factor receptor expression in clinical pain disorder tissues and role in neuronal sensitization. Pain Rep 2018;3:e676–e676.

[22] Elizur A, Adair-Kirk TL, Kelley DG, Griffin GL, Demello DE, Senior RM. Tumor necrosis factor-alpha from macrophages enhances LPS-induced clara cell expression of keratinocyte-derived chemokine. Am J Respir Cell Mol Biol 2008;38:8–15.

[23] Fang D, Kong L-Y, Cai J, Li S, Liu X-D, Han J-S, Xing G-G. Interleukin-6-mediated functional upregulation of TRPV1 receptors in dorsal root ganglion neurons through the activation of JAK/PI3K signaling pathway: roles in the development of bone cancer pain in a rat model. Pain 2015;156:1124–1144.

[24] Farr M, Garvey K, Bold AM, Kendall MJ, Bacon PA. Significance of the hydrogen ion concentration in synovial fluid in rheumatoid arthritis. Clin Exp Rheumatol 1985;3:99–104.

[25] Futami I, Ishijima M, Kaneko H, Tsuji K, Ichikawa-Tomikawa N, Sadatsuki R, Muneta T, Arikawa-Hirasawa E, Sekiya I, Kaneko K. Isolation and Characterization of Multipotential Mesenchymal Cells from the Mouse Synovium. PLoS ONE 2012;7:e45517.

[26] Gong W, Kolker SJ, Usachev Y, Walder RY, Boyle DL, Firestein GS, Sluka KA. Acid-sensing ion channel 3 decreases phosphorylation of extracellular signal-regulated kinases and induces synoviocyte cell death by increasing intracellular calcium. Arthritis Research & Therapy 2014;16:R121.

[27] Guo W, Yu D, Wang X, Luo C, Chen Y, Lei W, Wang C, Ge Y, Xue W, Tian Q, Gao X, Yao W. Anti-inflammatory effects of interleukin-23 receptor cytokine-binding homology region rebalance T cell distribution in rodent collagen-induced arthritis. Oncotarget 2016;7:31800–31813.

[28] Hannan MT. Epidemiologic perspectives on women and arthritis: An overview. Arthritis & Rheumatism 1996;9:424–434.

[29] Hardy RS, Hülso C, Liu Y, Gasparini SJ, Fong-Yee C, Tu J, Stoner S, Stewart PM, Raza K, Cooper MS, Seibel MJ, Zhou H. Characterisation of fibroblast-like synoviocytes from a murine model of joint inflammation. Arthritis Research & Therapy 2013;15:R24–R24.

[30] Hong R, Sur B, Yeom M, Lee B, Kim KS, Rodriguez JP, Lee S, Kang KS, Huh C-K, Lee SC, Hahm D-H. Anti-inflammatory and anti-arthritic effects of the ethanolic extract of Aralia continentalis Kitag. in IL-1β-stimulated human fibroblast-like synoviocytes and rodent models of polyarthritis and nociception. Phytomedicine 2018;38:45–56.

[31] Ita ME, Winkelstein BA. Concentration-Dependent Effects of Fibroblast-Like Synoviocytes on Collagen Gel Multiscale Biomechanics and Neuronal Signaling: Implications for Modeling Human Ligamentous Tissues. Journal of Biomechanical Engineering 2019;141. doi: 10.1115/1.4044051.

[32] Jones DS, Jenney AP, Swantek JL, Burke JM, Lauffenburger DA, Sorger PK. Profiling drugs for rheumatoid arthritis that inhibit synovial fibroblast activation. Nature Chemical Biology 2016;13:38.

[33] Jung H, Toth PT, White FA, Miller RJ. Monocyte chemoattractant protein-1 functions as a neuromodulator in dorsal root ganglia neurons. J Neurochem 2008;104:254–263.

[34] Kawashima M, Ogura N, Akutsu M, Ito K, Kondoh T. The anti-inflammatory effect of cyclooxygenase inhibitors in fibroblast-like synoviocytes from the human temporomandibular joint results from the suppression of PGE2 production. Journal of oral pathology & medicine: official publication of the International Association of Oral Pathologists and the American Academy of Oral Pathology 2013;42:499–506.

[35] Kochukov MY, McNearney TA, Yin H, Zhang L, Ma F, Ponomareva L, Abshire S, Westlund KN. Tumor Necrosis Factor-Alpha (TNF-α) Enhances Functional Thermal and Chemical Responses of TRP Cation Channels in Human Synoviocytes. Mol Pain 2009;5:1744-8069-5–49.

[36] Larsson S, Englund M, Struglics A, Lohmander LS. Interleukin-6 and tumor necrosis factor alpha in synovial fluid are associated with progression of radiographic knee osteoarthritis in subjects with previous meniscectomy. Osteoarthritis and Cartilage 2015;23:1906–1914.

[37] Lebre MC, Vieira PL, Tang MW, Aarrass S, Helder B, Newsom-Davis T, Tak PP, Screaton GR. Synovial IL-21/TNF-producing CD4+ T cells induce joint destruction in rheumatoid arthritis by inducing matrix metalloproteinase production by fibroblast-like synoviocytes. Journal of Leukocyte Biology 2017;101:775–783.

[38] Lee A, Qiao Y, Grigoriev G, Chen J, Park-Min K-H, Park SH, Ivashkiv LB, Kalliolias GD. Tumor necrosis factor α induces sustained signaling and a prolonged and unremitting inflammatory response in rheumatoid arthritis synovial fibroblasts. Arthritis Rheum 2013;65:928–938.

[39] Livak KJ, Schmittgen TD. Analysis of Relative Gene Expression Data Using Real-Time Quantitative PCR and the 2-ΔΔCT Method. Methods 2001;25:402–408.

[40] Llop-Guevara A, Porras M, Cendón C, Di Ceglie I, Siracusa F, Madarena F, Rinotas V, Gómez L, van Lent PL, Douni E, Chang HD, Kamradt T, Román J. Simultaneous inhibition of JAK and SYK kinases ameliorates chronic and destructive arthritis in mice. Arthritis Research & Therapy 2015;17:356.

[41] McCall MN, McMurray HR, Land H, Almudevar A. On non-detects in qPCR data. Bioinformatics 2014;30:2310–2316.

[42] Merabova N, Kaminski R, Krynska B, Amini S, Khalili K, Darbinyan A. JCV agnoprotein-induced reduction in CXCL5/LIX secretion by oligodendrocytes is associated with activation of apoptotic signaling in neurons. J Cell Physiol 2012;227:3119–3127.

[43] Miller RJ, Jung H, Bhangoo SK, White FA. Cytokine and chemokine regulation of sensory neuron function. Handb Exp Pharmacol 2009:417–449.

[44] Neumann E, Lefèvre S, Zimmermann B, Gay S, Müller-Ladner U. Rheumatoid arthritis progression mediated by activated synovial fibroblasts. Trends in Molecular Medicine 2010;16:458–468.

[45] Noss EH, Brenner MB. The role and therapeutic implications of fibroblast-like synoviocytes in inflammation and cartilage erosion in rheumatoid arthritis. Immunological Reviews 2008;223:252–270.

[46] Oh SB, Tran PB, Gillard SE, Hurley RW, Hammond DL, Miller RJ. Chemokines and Glycoprotein120 Produce Pain Hypersensitivity by Directly Exciting Primary Nociceptive Neurons. J Neurosci 2001;21:5027.

[47] Parsonage G, Falciani F, Burman A, Filer A, Ross E, Bofill M, Martin S, Salmon M, Buckley CD. Global gene expression profiles in fibroblasts from synovial, skin and lymphoid tissue reveals distinct cytokine and chemokine expression patterns. Thromb Haemost 2017;90:688–697.

[48] Picelli S, Faridani OR, Björklund ÅK, Winberg G, Sagasser S, Sandberg R. Full-length RNA-seq from single cells using Smart-seq2. Nature Protocols 2014;9:171–181.

[49] Rojewska E, Zychowska M, Piotrowska A, Kreiner G, Nalepa I, Mika J. Involvement of Macrophage Inflammatory Protein-1 Family Members in the Development of Diabetic Neuropathy and Their Contribution to Effectiveness of Morphine. Front Immunol 2018;9:494–494.

[50] Rosengren S, Boyle DL, Firestein GS. Acquisition, culture, and phenotyping of synovial fibroblasts. Methods in molecular medicine 2007;135:365–375.

[51] Safiri S, Kolahi AA, Hoy D, Smith E, Bettampadi D, Mansournia MA, Almasi-Hashiani A, Ashrafi-Asgarabad A, Moradi-Lakeh M, Qorbani M, Collins G, Woolf AD, March L, Cross M. Global, regional and national burden of rheumatoid arthritis 1990–2017: a systematic analysis of the Global Burden of Disease study 2017. Ann Rheum Dis 2019;78:1463.

[52] Sarria I, Ling J, Zhu MX, Gu JG. TRPM8 acute desensitization is mediated by calmodulin and requires PIP(2): distinction from tachyphylaxis. J Neurophysiol 2011;106:3056–3066.

[53] Saxne T, Palladino Jr MA, Heinegãrd D, Talal N, Wollheim FA. Detection of tumor necrosis factor α but not tumor necrosis factor β in rheumatoid arthritis synovial fluid and serum. Arthritis & Rheumatism 1988;31:1041–1045.

[54] Schaible H-G. Nociceptive neurons detect cytokines in arthritis. ARTHRITIS RESEARCH & THERAPY 2014;16.

[55] Scott BB, Weisbrot LM, Greenwood JD, Bogoch ER, Paige CJ, Keystone EC. Rheumatoid arthritis synovial fibroblast and U937 macrophage/monocyte cell line interaction in cartilage degradation. Arthritis & Rheumatism 1997;40:490–498.

[56] Sluka KA, Rasmussen LA, Edgar MM, O’Donnell JM, Walder RY, Kolker SJ, Boyle DL, Firestein GS. Acid-sensing ion channel 3 deficiency increases inflammation but decreases pain behavior in murine arthritis. 2013.

[57] Stefanini M, Martino C de, Zamboni L. Fixation of Ejaculated Spermatozoa for Electron Microscopy. Nature 1967;216:173–174.

[58] Syx D, Tran PB, Miller RE, Malfait A-M. Peripheral Mechanisms Contributing to Osteoarthritis Pain. Current Rheumatology Reports 2018;20:9.

[59] Taciak B, Białasek M, Braniewska A, Sas Z, Sawicka P, Kiraga Ł, Rygiel T, Król M. Evaluation of phenotypic and functional stability of RAW 264.7 cell line through serial passages. PLOS ONE 2018;13:e0198943.

[60] Taylor PC, Feldmann M. Anti-TNF biologic agents: still the therapy of choice for rheumatoid arthritis. Nature Reviews Rheumatology 2009;5:578.

[61] Tetta C, Camussi G, Modena V, Di Vittorio C, Baglioni C. Tumour necrosis factor in serum and synovial fluid of patients with active and severe rheumatoid arthritis. Ann Rheum Dis 1990;49:665–667.

[62] van Vollenhoven RF. Sex differences in rheumatoid arthritis: more than meets the eye.. BMC Med 2009;7:12–12.

[63] Wang YY, Chang RB, Waters HN, McKemy DD, Liman ER. The nociceptor ion channel TRPA1 is potentiated and inactivated by permeating calcium ions. J Biol Chem 2008;283:32691–32703.

[64] Westlund KN, Kochukov MY, Lu Y, McNearney TA. Impact of Central and Peripheral TRPV1 and ROS Levels on Proinflammatory Mediators and Nociceptive Behavior. Mol Pain 2010;6:1744-8069-6–46.

[65] Woolf CJ, Allchorne A, Safieh-Garabedian B, Poole S. Cytokines, nerve growth factor and inflammatory hyperalgesia: the contribution of tumour necrosis factor α. British Journal of Pharmacology 1997;121:417–424.

[66] Wright AJ, Husson ZMA, Hu D-E, Callejo G, Brindle KM, Smith EStJ. Increased hyperpolarized [1-13C] lactate production in a model of joint inflammation is not accompanied by tissue acidosis as assessed using hyperpolarized 13C-labelled bicarbonate. NMR in Biomedicine 2018:e3892-n/a.

[67] Yamamura Y, Gupta R, Morita Y, He X, Pai R, Endres J, Freiberg A, Chung K, Fox DA. Effector Function of Resting T Cells: Activation of Synovial Fibroblasts. J Immunol 2001;166:2270.

[68] Zhang F, Wei K, Slowikowski K, Fonseka CY, Rao DA, Kelly S, Goodman SM, Tabechian D, Hughes LB, Salomon-Escoto K, Watts GFM, Jonsson AH, Rangel-Moreno J, Meednu N, Rozo C, Apruzzese W, Eisenhaure TM, Lieb DJ, Boyle DL, Mandelin AM 2nd, Accelerating Medicines Partnership Rheumatoid Arthritis and Systemic Lupus Erythematosus (AMP RA/SLE) Consortium, Boyce BF, DiCarlo E, Gravallese EM, Gregersen PK, Moreland L, Firestein GS, Hacohen N, Nusbaum C, Lederer JA, Perlman H, Pitzalis C, Filer A, Holers VM, Bykerk VP, Donlin LT, Anolik JH, Brenner MB, Raychaudhuri S. Defining inflammatory cell states in rheumatoid arthritis joint synovial tissues by integrating single-cell transcriptomics and mass cytometry. Nat Immunol 2019;20:928–942.

